# Investigating the Contribution of Molecular-Enriched Functional Connectivity to Brain-Age Analysis

**DOI:** 10.1101/2025.10.16.682939

**Authors:** Marco Pinamonti, Manuela Moretto, Valentina Sammassimo, Marco Castellaro, Mattia Veronese

## Abstract

Brain-age prediction from neuroimaging data provides a proxy of biological aging, yet most models rely on structural magnetic resonance imaging (MRI), a modality that captures macroanatomy but offers limited biological specificity. We tested whether integrating molecular-enriched functional connectivity (FC), from resting-state functional MRI (rs-fMRI) data, improves brain-age prediction and biological explainability.

We analyzed MRI data of 2,120 healthy adults (1,243/877 F/M; 18–90 years) from three public datasets. Molecular-enriched connectivity maps were derived with Receptor-Enriched Analysis of functional Connectivity by Targets (REACT) using receptor-density templates for the dopamine (DAT), norepinephrine (NET), and serotonin (SERT) transporter systems. Support vector regression models were applied to predict chronological age from molecular-enriched FC, structural morphometry, or both combined. The effect of multi-site variability was mitigated via ComBat harmonization with and without Empirical Bayes pooling. We additionally conducted a common-parcellation analysis to assess the impact of differing parcellations between modalities.

Single-transporter molecular-enriched FC explained up to 51% of age variance. The most predictive transporter varied by dataset, with DAT dominating in the harmonized and common-parcellation settings. Combining the three molecular-enriched maps consistently improved prediction over any single map and increased explained variance up to 64%. In the merged multi-site cohort using a common parcellation, augmenting structural information with transporter-enriched FC reduced mean absolute error (MAE) from 6.02 to 5.81 years, supporting complementarity of the two modalities. In contrast, when different parcellations were applied, incorporating molecular-enriched FC into brain age prediction resulted in a 2% higher MAE compared to structural morphometry alone, suggesting that parcellation mismatch may obscure the functional contributions.

In conclusion, molecular-enriched FC is a feasible and biologically informative extension to brain-age modeling, enhancing prediction and interpretability with respect to neurotransmitter systems.

## 1 Introduction

Worldwide, life expectancy has risen dramatically over recent decades; demographic forecasts indicate that by 2050, approximately 16% of the global population will be older than 65 years, nearly double the 9% recorded in 2019 (United Nations, Department of Economic and Social Affairs, Population Division, 2019). This demographic shift is already imposing major challenges for sustainability of healthcare systems and social services, as older age is accompanied by a higher burden of chronic diseases and disabilities (World Health Organization, 2024). The clinical and public health implications are urgent and far-reaching, reflecting a rapidly growing global burden and escalating economic costs that motivate sensitive markers of aging and early risk stratification (Alzheimer’s Disease International, 2024; Livingston et al., 2024).

Aging is a complex physiological process characterized by the gradual accumulation of molecular and cellular damage, leading to functional decline and increased vulnerability to disease (Blinkouskaya et al., 2021). Advanced age represents the greatest risk factor for numerous chronic conditions, including cancers, cardiovascular disease, and neurodegenerative disorders such as dementia and Alzheimer’s disease (MacDonald & Pike, 2021). Yet the rate at which aging unfolds varies markedly among individuals, reflecting a dynamic interplay of genetic, lifestyle, and environmental influences (Moskalev, 2019; Udeh-Momoh et al., 2025). Consequently, chronological age often fails to capture an individual’s biological state. The concept of biological age has therefore gained traction as a more informative metric across organ systems, with molecular and phenotypic clocks used to stratify risk and to monitor intervention effects (Moskalev, 2019).

Within this framework, neuroimaging has been widely used to estimate the biological age of the brain, which is thought to reflect brain-specific aging processes (Cole & Franke, 2017; Franke & Gaser, 2019). Often referred to as “brain-age”, this framework consists of training machine-learning models on large normative magnetic resonance imaging (MRI) datasets of T1-weighted structural images to predict chronological age (Cole et al., 2015; Franke et al., 2010). Hence for a new individual, the brain-age gap (sometimes also referred to as the brain-age Δ) can be calculated as the difference between the predicted brain-age and the person’s actual chronological age. A positive Δ indicates an older-appearing brain than expected for chronological age, whereas a negative Δ indicates a younger-appearing brain. In applications, Δ can be summarized for group comparisons (e.g., patients vs. controls) or tracked longitudinally within individuals for risk stratification and to gauge intervention effects. This metric is typically interpreted as an index of accelerated or decelerated brain aging: large positive Δ is associated with diverse disorders, including schizophrenia (Koutsouleris et al., 2013; Nenadić et al., 2017; Schnack et al., 2016), dementia (Wang et al., 2019), Alzheimer’s disease (Franke & Gaser, 2012), multiple sclerosis (Høgestøl et al., 2019), epilepsy (Pardoe et al., 2017), HIV infection (Cole, Underwood, et al., 2017), Down syndrome (Cole, Annus, et al., 2017), and type 2 diabetes (Franke et al., 2013). On the contrary, healthy habits in terms of diet and environment have been shown to slow the aging process in animal models, resulting in negative Δ (Brusini et al., 2022).

To enhance the sensitivity and accuracy of brain-age models, functional connectivity (FC) features derived from resting-state functional MRI (rs-fMRI) have been incorporated (Liem et al., 2017). This follows evidence from numerous studies showing that normal aging is characterized by systematic alterations in network coherence and inter-network communication, assessed using both static (Betzel et al., 2014; Ferreira & Busatto, 2013) and dynamic approaches (Chen et al., 2019; Moretto et al., 2022; Tian et al., 2018; Xia et al., 2019). On the basis of these age-related alterations in resting-state FC (rs-FC) features, a few studies have attempted to predict chronological age using machine-learning models trained on rs-FC patterns (Bi et al., 2024; Millar et al., 2022). In Bi et al. (2024), age was predicted with a mean absolute error (MAE) of 6–7 years, with predictions explaining approximately 74% of age variability, approaching but not surpassing the precision of structural MRI–based models that stand at 93% (Cole et al., 2019; Gutiérrez Becker et al., 2018). Notably, in preclinical Alzheimer’s disease, deviations in FC-based brain age have been detected even before overt memory symptoms emerge (Millar et al., 2022), suggesting that rs-FC may capture early functional changes missed by structural imaging.

The BOLD signal reflects changes in the local balance of oxyand deoxyhemoglobin arising from neurovascular coupling and thus provides an indirect hemodynamic proxy for neural activity. Consequently, despite its potential for studying brain age, rs-fMRI lacks neurochemical specificity (Logothetis, 2008). To address this limitation, Dipasquale et al. (2019) introduced the Receptor-Enriched Analysis of functional Connectivity by Targets (REACT), which integrates BOLD fluctuations with normative receptor-density maps derived from PET or SPECT, yielding molecular-enriched functional connectivity that incorporates neurochemical information. After its introduction, REACT has been increasingly applied in diverse contexts, including pharmacological challenges such as MDMA (Dipasquale et al., 2019; Singleton et al., 2023), LSD (Lawn et al., 2022), and methylphenidate (Dipasquale et al., 2020); neurological conditions such as multiple sclerosis (Cercignani et al., 2021), Parkinson’s disease (Di Vico & Moretto et al., 2025), and visual snow syndrome and migraine with aura (Puledda et al., 2023); as well as in chronic pain (Martins et al., 2021). Moreover, by integrating REACT with normative modeling across the healthy lifespan, Lawn et al. (2024) showed that molecular-enriched network deviations systematically track age-related functional reorganization, providing a sensitive molecularly informed biomarker of brain-age (Lawn et al., 2024). Integrating such molecularly informed functional features with established structural metrics therefore holds promise for refining brain-age estimates and elucidating the neurochemical substrates of age-related brain changes.

Building molecular-enriched rs-fMRI into brain-age paradigms comes at the cost of a few methodological challenges. First and foremost, cohorts must be large enough (N>2,000) to match the sample sizes that have made structure-only models successful (Han et al., 2020; Varoquaux, 2018; Yu et al., 2024). Because single datasets rarely reach this scale, assembling an adequately powered cohort requires aggregating data across sites, scanners, and protocols. Such aggregation demands principled harmonization to attenuate non-biological site effects while preserving age-related signal. In parallel, study design should support generalizability by employing broad normative training samples, enforcing consistent parcellation across modalities, and evaluating performance across sites and acquisition conditions.

Motivated by the need for biologically informative features, the present study investigates whether applying REACT to integrate neurochemical information into rs-fMRI data from 2,120 healthy adults might improve brain-age prediction. Specifically, it aims to (i) quantify the standalone prediction accuracy of molecular-enriched FC and benchmark it against structural T1-weighted (T1w) MRI — evaluating each modality, both separately and combined, under the hypothesis that molecular-enriched FC provides complementary variance that enhances overall accuracy — and (ii) assess the multi-site problem by comparing alternative harmonization strategies for mitigating scanner and site effects across structural morphometry, molecular-enriched FC, and receptor-density atlas features within a unified modeling pipeline.

## 2 Methods

### 2.1 Ethics statement

This study analyzed de-identified human neuroimaging and phenotypic data from three publicly available adult cohorts. Each source study obtained approval from its local research ethics committee/institutional review board, and all participants provided written informed consent for data collection and for data sharing. Data access and use complied with each resource’s data-use terms; no attempts at reidentification were made, and all reporting is at the group level. No new data were collected, and no animal research was involved. Receptor-density templates used in REACT were taken from previously published PET/SPECT atlases that were acquired under the oversight of their originating studies’ ethics committees and reused under their licenses. All procedures adhered to the Declaration of Helsinki and applicable data-protection regulations.

### 2.2 Datasets

This study assembled a large cohort of normative individuals with both rs-fMRI and T1w MRI by pooling three publicly available lifespan studies focused on aging: the “Cambridge Centre for Ageing and Neuroscience” study (Cam-CAN) (Shafto et al., 2014; Taylor et al., 2017), the “Lifespan Human Connectome Project in Aging” (HCP-Aging) (Bookheimer et al., 2019; Harms et al., 2018), and the “Nathan Kline Institute Rockland Sample” (NKI-RS) (Nooner et al., 2012).

Across these cohorts, the “healthy” status of participants was defined pragmatically for normative aging research as community-dwelling adults who could provide informed consent and safely undergo MRI. Exclusions focused on major diagnosed disease across neurological (e.g., stroke, Parkinson’s disease, epilepsy), psychiatric (e.g., schizophrenia, bipolar disorder), and systemic medical domains (e.g., active cancer therapy, uncontrolled cardiovascular disease), as well as cognitive impairment, current substance misuse, and MRI contraindications; common, stable age-related conditions (e.g., well-controlled hypertension) were generally permitted. Cam-CAN and HCP-Aging explicitly screened for major neurological and psychiatric illness within this “typical aging” framework, whereas NKI-RS used community-ascertained recruitment calibrated to the age, sex, ethnicity, and socioeconomic profile of Rockland County, New York, to approximate broader U.S. demographics (Bookheimer et al., 2019; Nooner et al., 2012; Shafto et al., 2014).

From these datasets, we included only participants with good quality T1w and rs-fMRI data resulting in 653 individuals from Cam-CAN, 724 from HCP-Aging and 978 from NKI-RS. For all these studies, imaging was acquired on 3 T Siemens systems with 32-channel head coils (TIM Trio for Cam-CAN and NKI-RS, Prisma for HCP-Aging). The multi-site study (HCP-Aging) implemented harmonized, site-matched sequences across four centers; Cam-CAN and NKI-RS were collected at single sites using consistent protocols. Sequence parameters are summarized in Table 1 (structural) and Table 2 (functional).

**Table 1:**
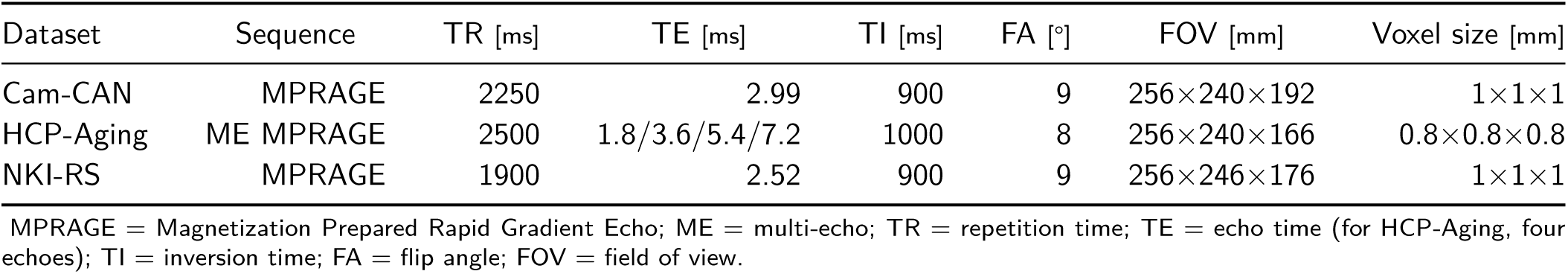
T1-weighted structural MRI acquisition parameters for each cohort.

**Table 2:** Resting–state functional MRI acquisition parameters for each cohort.

| Dataset | Sequence | TR [ms] | TE [ms] | FA [°] | FOV [mm] | Voxel size [mm] | Slices | Volumes | Time [min:sec] |
| --- | --- | --- | --- | --- | --- | --- | --- | --- | --- |
| Cam-CAN | SB GE-EPI | 1970 | 30 | 78 | 192×192 | 3×3×4.4 | 32 | 261 | 8:34 |
| HCP-Aging | MB8 GE-EPI | 870 | 37 | 52 | 208×208 | 2×2×2 | 72 | 488 | 7:05 |
| NKI-RS | MB4 GE-EPI | 1400 | 30 | 65 | 224×224 | 2×2×2 | 64 | 418 | 9:45 |

SB = single-band; MB = multi-band; GE = gradient-echo; EPI = echo-planar imaging; TR = repetition time; TE = echo time; TI = inversion time; FA = flip angle; FOV = field of view.

### 2.3 Image analysis

Across cohorts, we obtained raw T1w and rs-fMRI images and applied a uniform preprocessing workflow to ensure cross-cohort comparability. Raw images were organized according to Brain Imaging Data Structure (BIDS) specifications (Gorgolewski et al., 2016) and then processed with standardized BIDS Apps (BIDS Apps Community, 2025). Each dataset was then checked with the bids-validator (v1.14.6)^1^, which confirmed full compliance with the specification (Blair et al., 2024).

#### 2.3.1 Quality control of raw images

The large number of scans downloaded from Cam-CAN, HCP-Aging, and NKI-RS precluded exhaustive manual inspection; therefore, image quality was assessed with MRIQC^2^ (v24.1.0) (Esteban et al., 2017). MRIQC has become a standard tool for automated QC of structural and functional MRI data (Botvinik-Nezer et al., 2019; Nakua et al., 2023; Nitsch et al., 2024; Provins et al., 2023). The tool was run independently for each cohort and modality, and the resulting group-level reports were inspected. Any scan with image-quality metrics (IQMs) outside the inner Tukey fences (i.e., below *Q*_1_ − 1.5, IQR or above *Q*_3_ + 1.5, IQR) was flagged for review according to the non-parametric outlier rule (Tukey, 1977). We focused on IQMs that are particularly sensitive to motion and other artifacts: for T1w images, signal-to-noise ratio (SNR), gray–white matter contrast-to-noise ratio, and the entropy-focus criterion; for rs-fMRI runs, temporal SNR, mean framewise displacement, and temporal derivative of the variance of the BOLD signal (DVARS). All flagged scans were visually inspected; scans exhibiting motion, ringing, cropping or other obvious defects were excluded, and the remaining data were processed further.

#### 2.3.2 Preprocessing

T1w and rs-fMRI images were processed with a three-stage workflow that combined fMRIPrep (v23.2.2), XCP-D (v0.7.4), and FSL (v7.3.2). fMRIPrep is an automated, BIDS-compatible pipeline that delivers analysis-ready structural and functional derivatives by integrating core routines from Advanced Normalization Tools (ANTs), FMRIB Software Library (FSL), FreeSurfer (FS), and Analysis of Functional NeuroImages (AFNI) (Esteban et al., 2019, 2020); XCP-D is a post-processing toolkit that implements state-of-the-art nuisance regression and connectivity derivatives (Mehta et al., 2024); and FSL is a long-standing MRI toolbox offering spatial filtering, registration, and statistical modeling (Jenkinson et al., 2012; Smith et al., 2004). Together, these tools constitute one of the most widely adopted preprocessing stacks in contemporary neuroimaging studies (Bowring et al., 2019; Dugré et al., 2025).

##### Structural and functional preprocessing with fMRIPrep

Each participant’s T1w image was corrected for intensity non-uniformity with N4 bias field correction (ANTs 2.5.0) (Tustison et al., 2010), then skull-stripped via the Nipype implementation of antsBrainExtraction.sh using the OASIS30ANTs target (TemplateFlow v23.1.0) (Ciric et al., 2022). Tissue segmentation into cerebrospinal fluid (CSF), white matter (WM), and gray matter (GM) was performed on the brain-extracted T1w with fast (FSL 7.3.2) (Zhang et al., 2001). Cortical surfaces were reconstructed with recon-all (FreeSurfer 7.3.2) (Fischl, 2012), and the initial brain mask was refined by reconciling ANTs- and FreeSurfer-derived GM segmentations via a Mindboggle-based algorithm (Klein & Tourville, 2012). Finally, the T1w was nonlinearly normalized to the MNI ICBM 152 NL-6 Asym template (2 mm isotropic) using antsRegistration (ANTs 2.5.0) through TemplateFlow (Fonov et al., 2011).

For each rs-fMRI run, a reference volume was generated for head-motion correction, and motion parameters were estimated with mcflirt (FSL) before any filtering (Jenkinson et al., 2012). The BOLD reference was coregistered to the T1w via boundary-based registration (bbregister, FreeSurfer) (Greve & Fischl, 2009), with six degrees of freedom. All spatial transformations (head-motion, BOLD to T1w, T1w to MNI) were composed and applied in a single interpolation step using nitransforms with cubic B-spline interpolation.

From fMRIPrep we retained only: (i) spatial transforms and the preprocessed BOLD in native and MNI spaces, and (ii) the _confounds.tsv columns needed to identify motion-affected frames (framewise displacement (FD) and DVARS) plus six rigid-body parameters (with their temporal derivatives and quadratics). Volumes with FD>0.5 mm or DVARS>1.5 *σ* were flagged for interpolation downstream (Power et al., 2012); no temporal filtering or nuisance regression was performed at this stage.

##### Post-processing with XCP-D

The eXtensible Connectivity Pipeline for DCAN (XCP-D) was used to post-process fMRIPrep derivatives (Gorgolewski et al., 2011; Mehta et al., 2024). Confound regression followed the “acompcor” nuisance model, defined as six motion parameters plus their temporal derivatives and the top five aCompCor components from WM and CSF (Behzadi et al., 2007; Satterthwaite et al., 2013). Frames flagged as high-motion were cubic-spline interpolated; both data and confounds were then high-pass filtered (second-order Butterworth, 0.001 Hz cutoff) and denoised via linear regression of the nuisance set and cosine bases (Ciric et al., 2018). Because downstream analyses required continuous time series, we retained the interpolated, non-censored residual BOLD image produced by XCP-D (no spatial smoothing in this step).

##### Spatial smoothing with FSL

The interpolated residual BOLD images were spatially smoothed in FSL 7.3.2 with a Gaussian kernel with an FWHM of 6 mm. These smoothed, denoised BOLD volumes were used for all subsequent FC and brain-age analyses.

### 2.4 Molecular-enriched functional connectivity

Using the preprocessed BOLD time series as input, we derived molecular-enriched FC maps that fuse neurotransmitter-specific information from positron-emission tomography (PET)/single-photon emission computed tomography (SPECT) templates with rs-fMRI, thus augmenting the hemodynamic proxy of neuronal activity with biologically grounded molecular context.

#### 2.4.1 Receptor-Enriched Analysis of Functional Connectivity by Targets (REACT)

The REACT framework enriches rs-fMRI BOLD fluctuations with molecular density data derived from publicly available PET/SPECT templates to derive neurotransmitter-informed FC maps (Dipasquale et al., 2019). It proceeds in two steps as depicted in Figure 1: a) spatial regression: each preprocessed rs-fMRI run is regressed in time onto a receptor-density template, yielding a subject- and target-specific weight time series that quantifies how strongly that molecular distribution modulates ongoing BOLD fluctuations; b) temporal regression: this weight time series is then entered as a regressor in space against the voxel-wise BOLD signal, producing a voxel-wise connectivity map enriched for the same receptor target, and thus revealing neurochemically informed circuit fingerprints.

**Figure 1:**
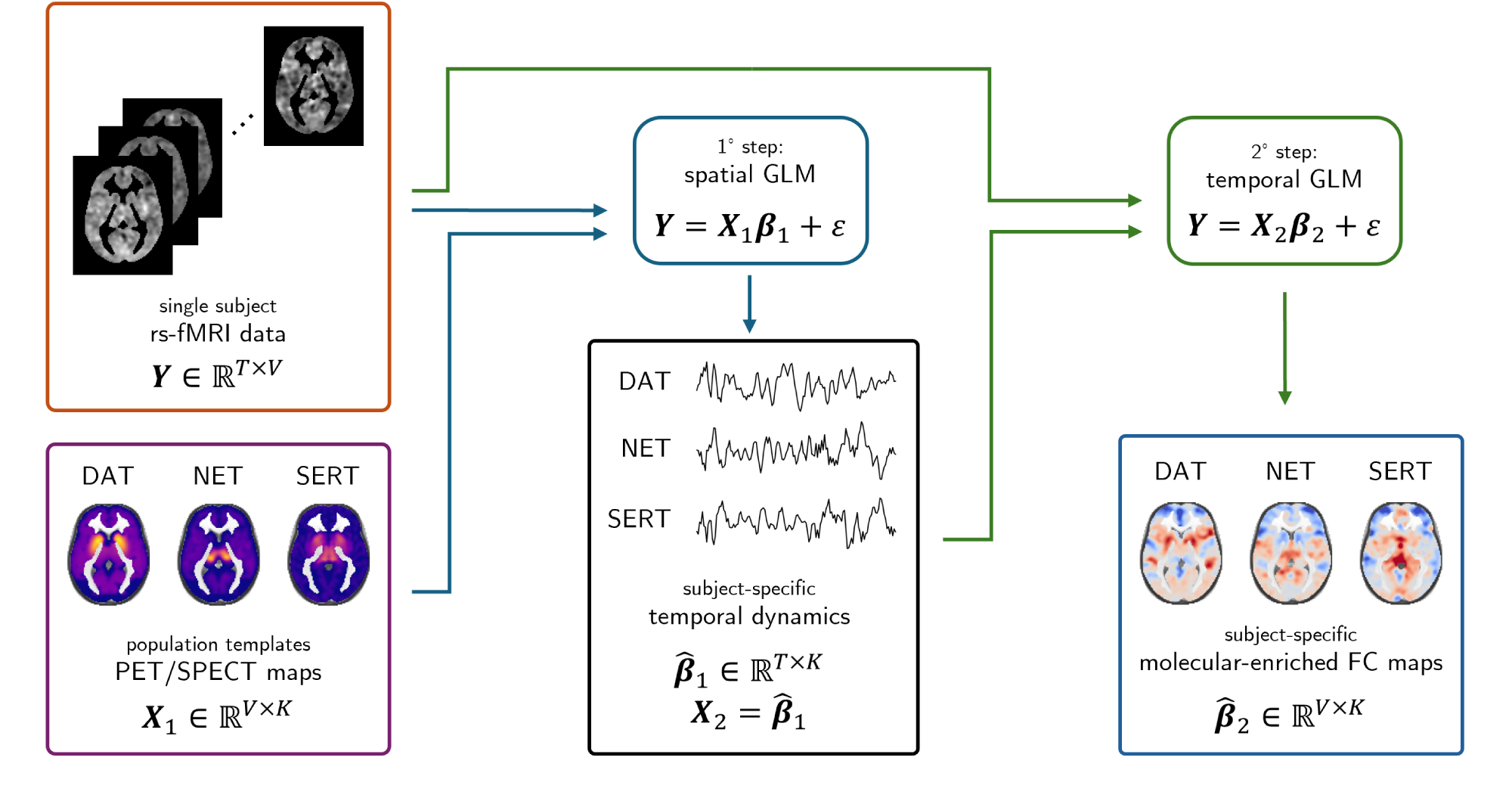
REACT workflow. The REACT framework derives neurotransmitter-informed FC from rs-fMRI by combining subject BOLD data with population PET/SPECT receptor-density templates. Inputs are the BOLD matrix *Y* ∈ ℝ*^T^ ^×V^* and the template matrix *X*_1_ ∈ ℝ*^V^ ^×K^* (here, DAT, NET, and SERT). *Step 1 - spatial GLM: Y* = *X*_1_*β*_1_ + *ε* estimates, for each target, a subject-specific weight time series 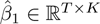 indexing how strongly that molecular distribution modulates ongoing BOLD fluctuations. *Step 2 - temporal GLM:* setting 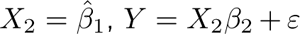 yields voxel-wise maps 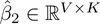 that represent receptor-enriched connectivity patterns.

#### 2.4.2 Neuroreceptor templates

Three templates for the transporters of dopamine (DAT), norepinephrine (NET), and serotonin (SERT) were selected. The DAT map was constructed from [^123^I]-ioflupane SPECT images acquired in 30 neurologically healthy adults showing no signs of nigrostriatal impairment (García-Gómez et al., 2013). The NET template was generated by averaging individual [^11^C]-MRB PET parametric maps from a cohort of 10 healthy subjects (Hesse et al., 2017). The SERT template was derived by pooling [^11^C]-DASB PET images from 210 healthy volunteers to generate a high-resolution binding distribution (Beliveau et al., 2017). All three templates were normalized into MNI152NLin6Asym space with 2 mm isotropic voxels, and scaled between 0 and 1.

These targets were selected a priori because monoaminergic transporters exhibit well-characterized age-related alterations and regulate large-scale network dynamics detectable with BOLD functional connectivity (Buchert et al., 2006; Ding et al., 2010; Kanel et al., 2023; Karrer et al., 2017). Dopaminergic markers, including transporter and receptor availability, decline across adulthood and relate to individual differences in cognitive performance (Karrer et al., 2017). Noradrenergic integrity, encompassing the locus coeruleus and its cortical projections, shows age-related change with established links to attention and arousal, and PET studies indicate reduced NET binding with age (Ding et al., 2010; Mather & Harley, 2016). Serotonergic terminals and transporter availability also display regional age-related reductions (Buchert et al., 2006; Kanel et al., 2023). All together, DAT, NET, and SERT provide biologically interpretable axes for molecular-enriched FC in brain-age models (Hansen et al., 2022). Although additional neurotransmitter systems and receptor subtypes also show age-dependent variation, we restricted the present analysis to monoaminergic transporters for methodological and comparability reasons: first, to maintain continuity with prior REACT applications and second, to limit spatial collinearity among molecular maps, particularly that arising between receptor subtypes with overlapping distributions, which can destabilize the two-stage regression and hinder interpretability (Beliveau et al., 2017; Dipasquale et al., 2019; Hansen et al., 2022).

#### 2.4.3 REACT implementation

The react-fmri tool (https://github.com/ottaviadipasquale/react-fmri) requires two binary masks for its two-stage regression: i) a Stage-1 mask restricting the temporal regression to voxels common to each transporter template and to cortical gray matter (GM); ii) a Stage-2 mask for the spatial regression, here taken as the GM mask alone. We generated the Stage-1 mask as the intersection between each transporter template and the generic GM mask supplied with REACT, and the Stage-2 mask as that GM mask alone.

Processing then proceeded in two steps. i) Each participant’s preprocessed BOLD data were processed with the command-line utility react-fmri, once per transporter template, yielding subject- and target-specific weight images and transporter-enriched functional-connectivity maps in MNI152NLin6Asym space. ii) Every transporter-enriched connectivity map was parcellated into 247 regions of interest (ROIs) — 200 cortical parcels from the Schaefer 7-network atlas (Schaefer et al., 2017), 15 subcortical regions from the Harvard–Oxford structural atlas (FMRIB Analysis Group, University of Oxford & Harvard Center for Morphometric Analysis, 2023; Smith et al., 2004), and 32 cerebellar regions from the SUIT atlas (Diedrichsen, 2009) — and the resulting #subjects×247 feature matrices (one per transporter) were exported as tab-separated value files for downstream brain-age modeling.

### 2.5 Brain-age modeling workflow

The preprocessing stage yielded, for every dataset and every transmitter system, a #subjects×247 feature matrix that served as input to a machine-learning pipeline for estimating brain-age.

#### Feature blocks and models comparison

Across all analyses we considered six model configurations: three transmitter-specific models (DAT, NET, SERT), each using 247 molecular-enriched FC regional features (mean receptor-enriched connectivity within each of the 247 ROIs defined above); a concatenated molecular-enriched features (MEF) block that stacks the three transmitter sets for a total of 3×247=741 features; a structural morphometry block (SMF, 649 features) comprising (a) 85 subcortical/global volumetric indices (Fischl et al., 2002), (b) 2×31 cortical regions × 8 metrics (GM volume, cortical surface area, mean cortical thickness, thickness standard deviation, folding index, mean curvature, Gaussian curvature, curvature index) from the Desikan–Killiany-Tourville atlas (DKT) parcellation (Desikan et al., 2006), and (c) 68 regional WM volumes; and a multimodal union (MMF) obtained by concatenating MEF and SMF for a total of 741+649=1,390 features.

#### Learning algorithm

We used support vector regression (SVR) with a radial-basis-function kernel as a compromise between predictive accuracy and computational efficiency, consistent with earlier bench-marking (Pinamonti & Sammassimo et al., 2025). All analyses were executed in a Conda environment on a Slurm-managed cluster to ensure software consistency. Models predicted chronological age from the feature blocks above and were evaluated with 10-fold cross-validation repeated 10 times (folds stratified by age). Within each split, all predictors were scaled with RobustScaler (median/IQR; scikit-learn), fitted on the training fold only and applied to the test data. Hyper-parameters (C, *γ*, *ε*) were tuned by nested grid search within each training fold.

To structure the study, we organized three complementary analysis blocks: (i) *single-cohort models* to test whether molecular-enriched FC improves brain-age estimation within individual datasets; (ii) *merged-cohort models* to achieve sample sizes typical of structural brain-age work and to assess whether multi-site harmonization is required and what are its optimal settings; (iii) a *common-parcellation analysis* to isolate parcellation as a potential confound by aligning structural and functional features on the same atlas.

#### 2.5.1 Single-cohort models: testing the added value of molecular-enriched FC

To test whether molecular-enriched FC added predictive information for brain-age estimation, we evaluated six pre-specified model configurations within each dataset separately. These configurations spanned structure-only models, neurotransmitter-specific FC variants, and structure+functional combinations, all trained and validated under the same preprocessing, scaling, and nested cross-validation procedures described above.

#### 2.5.2 Merged-cohort models: assessing harmonization for multi-site integration

To determine whether batch correction was necessary and how best to implement it, we merged Cam-CAN, HCP-Aging, and NKI-RS to reach sample sizes typical of structural brain-age studies and compared models before and after harmonization.

We first trained models on non-harmonized data to quantify multi-site effects, and then repeated the analyses after neuroCombat correction (with and without empirical Bayes (EB)), using acquisition site as batch and preserving age and sex as covariates (Fortin et al., 2018). Two harmonization configurations were tested: (a) per-feature location/scale adjustment *without* EB pooling (no information borrowing across features), and (b) ComBat *with* EB pooling (Fortin et al., 2017; Johnson et al., 2007). For the latter, rather than estimating a single set of ComBat parameters across all features jointly, we first partitioned the merged feature matrix into homogeneous “subgroups” to respect measurement type and biological domain: for FreeSurfer structural features, we grouped variables derived from the same .stats file and of the same nature (volumes, surface areas, cortical folding or curvature); for functional connectivity, we grouped edges by neuroreceptor system (DAT, NET, SERT) and by a single atlas-defined parcel set (one of the seven Schaefer networks, the combined subcortical mask, or the cerebellar mask). We then ran neuroCombat separately on each subgroup, with the argument eb=True to invoke EB pooling for that subgroup only. This strategy fits distinct empirical-Bayes priors and hyper-parameters for each feature domain, so that borrowing of strength occurs only among features of the same type, preserving modality-specific biological variance while attenuating scanner-and site-related effects within each subgroup (Fortin et al., 2017; Johnson et al., 2007). After harmonization, the corrected sub-matrices were recombined for downstream cross-validation and SVR modeling. For every harmonization setting, we retrained transmitter-specific models, the concatenated functional model, the FreeSurfer structural model, and the full multimodal model, following the same cross-validation protocol.

#### 2.5.3 Common-parcellation analysis: testing parcellation effects on multimodal gains

To isolate atlas mismatch as a potential confound in the merged-cohort setting, we projected both structural and neurotransmitter-weighted FC features onto identical subject-specific DKT parcels and retrained transmitter-specific, structural, and combined models after neuroCombat harmonization (no EB). This analysis tested whether molecular-enriched FC contributed information beyond FreeSurfer-derived structure when both were summarized on the same atlas.

#### 2.5.4 Evaluation metrics

Model accuracy was quantified with five complementary statistics computed in every test fold, following the evaluation framework proposed by Baecker et al. (2021). MAE measured the average unsigned deviation between predicted and chronological age, whereas root-mean-squared error (RMSE) penalized larger residuals by squaring them before averaging and taking the square root. Pearson’s correlation coefficient (*r*) captured the linear association between true and estimated ages, and the prediction coefficient of determination (*R*^2^) summarized the proportion of age variance explained in unseen data. Bias was assessed as Spearman’s rank correlation (*ρ*) between chronological age and the prediction error 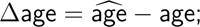; values near zero indicate that errors are not systematically age-dependent.

## 3 Results

### 3.1 Datasets

Following MRIQC screening, motion censoring, and preprocessing, we excluded 226 participants, leaving 2,128 adults eligible for analysis across the three source studies. Because the HCP-Aging cohort contained only eight participants older than 90 years, too few for robust model estimation at that extreme age range, we excluded those individuals. The final analytic sample therefore comprised 2,120 participants. Table 3 summarizes sample size, sex distribution, and age statistics for each dataset. Further information on participant exclusion after MRIQC and preprocessing is reported in the Supplementary Materials.

**Table 3:** Demographic characteristics of the quality-controlled cohorts.

| Dataset | *N* | Female / Male | Age, mean ± SD [years] | Age range [years] |
| --- | --- | --- | --- | --- |
| Cam-CAN | 627 | 315 / 312 | 53.8 ± 18.5 | 18–88 |
| HCP-Aging | 599 | 337 / 262 | 58.6 ± 14.7 | 36–90 |
| NKI-RS | 894 | 591 / 303 | 46.7 ± 17.7 | 18–85 |
| **Total** | **2,120** | **1,243 / 877** | 52.2 ± 17.9 | 18–90 |

### 3.2 Qualitative characteristics of the molecular-enriched FC maps

Figure 2 displays the group-mean transporter-enriched FC maps for each dataset (Cam-CAN, HCP-Aging, NKI-RS) and transmitter target (DAT, NET, SERT). Across cohorts, peak connectivity localizes to neuroanatomically plausible loci: the DAT map emphasizes the anterior striatum and caudate head; the NET map highlights the cerebellar vermis-brainstem junction; and the SERT map shows maximal signal in the midline cerebellar dentate/brainstem region. These spatial patterns align with prior reports of monoamine transporter–enriched connectivity (Cercignani et al., 2021; Coray et al., 2023; Dipasquale et al., 2023).

**Figure 2:**
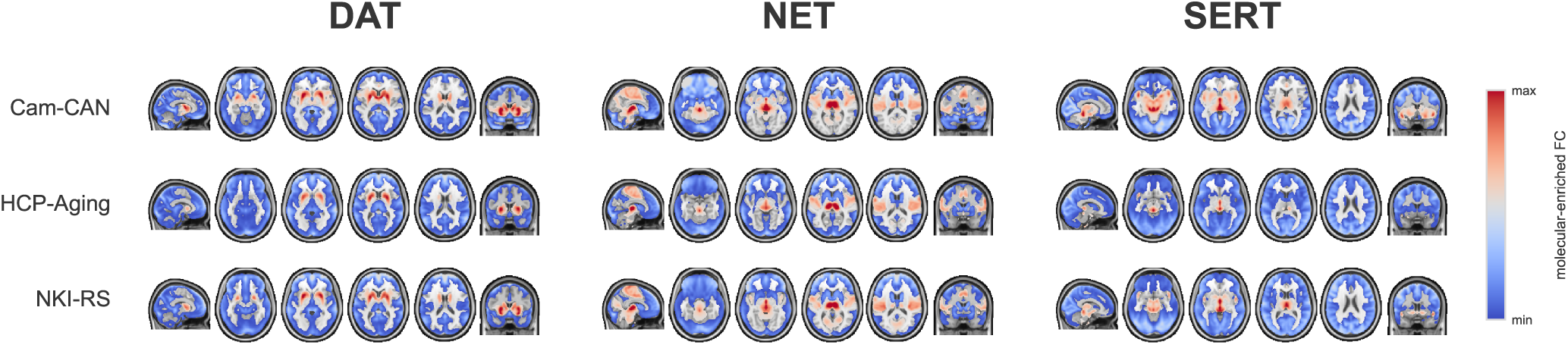
Group-mean molecular-enriched FC maps derived with REACT for the dopamine (DAT), norepinephrine (NET), and serotonin (SERT) transporters. Rows correspond to datasets (Cam–CAN, HCP–Aging, NKI–RS), columns to transporter targets. The color scale indicates relative connectivity (arbitrary units; min–max as shown). Consistent across datasets, DAT emphasizes anterior striatum/caudate head, NET emphasizes the cerebellar vermis–brainstem junction, and SERT emphasizes the midline cerebellar dentate/brainstem region, matching previously reported topographies (Cercignani et al., 2021; Coray et al., 2023; Dipasquale et al., 2023).

### 3.3 Brain-age modeling

The sections below report prediction accuracy and error metrics for the six model configurations including three individual neuroreceptor-enriched models, their combined model MEF, the structural model SMF and the multimodal union MMF.

#### 3.3.1 Single-cohort models: molecular-enriched FC generally adds predictive value

Table 4 first lists the accuracy obtained from the single transporter maps. Across cohorts, the DAT, NET, and SERT features yielded moderate explained variance in chronological age (*R*^2^ of 0.50–0.52 in Cam–CAN, 0.27–0.40 in HCP-Aging, 0.38–0.45 in NKI–RS), with MAEs clustered around 9.27–11.43 years. When the three functional maps were concatenated (MEF model), performance improved consistently, reducing the typical MAE by 0.96–1.82 years relative to the best single map and raising *R*^2^ into the 0.50–0.64 range. FreeSurfer-derived structural metrics (SMF model) remained the strongest single modality, posting the lowest MAEs (5.55–5.76 years) and the highest coefficients of determination (*R*^2^ of 0.78–0.85). Merging MEF with SMF (MMF model) retained this high accuracy and slightly surpassed pure structure in HCP-Aging (MAE = 5.49 vs. 5.55) and NKI-RS (MAE = 5.68 vs. 5.76). In summary, FS-derived structural features were most accurate in Cam–CAN, whereas the combined structural+functional set delivered the best result in HCP-Aging and NKI–RS. As illustrated in Figure 3, these numeric differences are echoed graphically: panels (a)–(b) show that MAE falls and *R*^2^ rises from the single-transporter maps (DAT, NET, SERT) to the concatenated REACT features, and again to the structural and multimodal models.

**Table 4:**
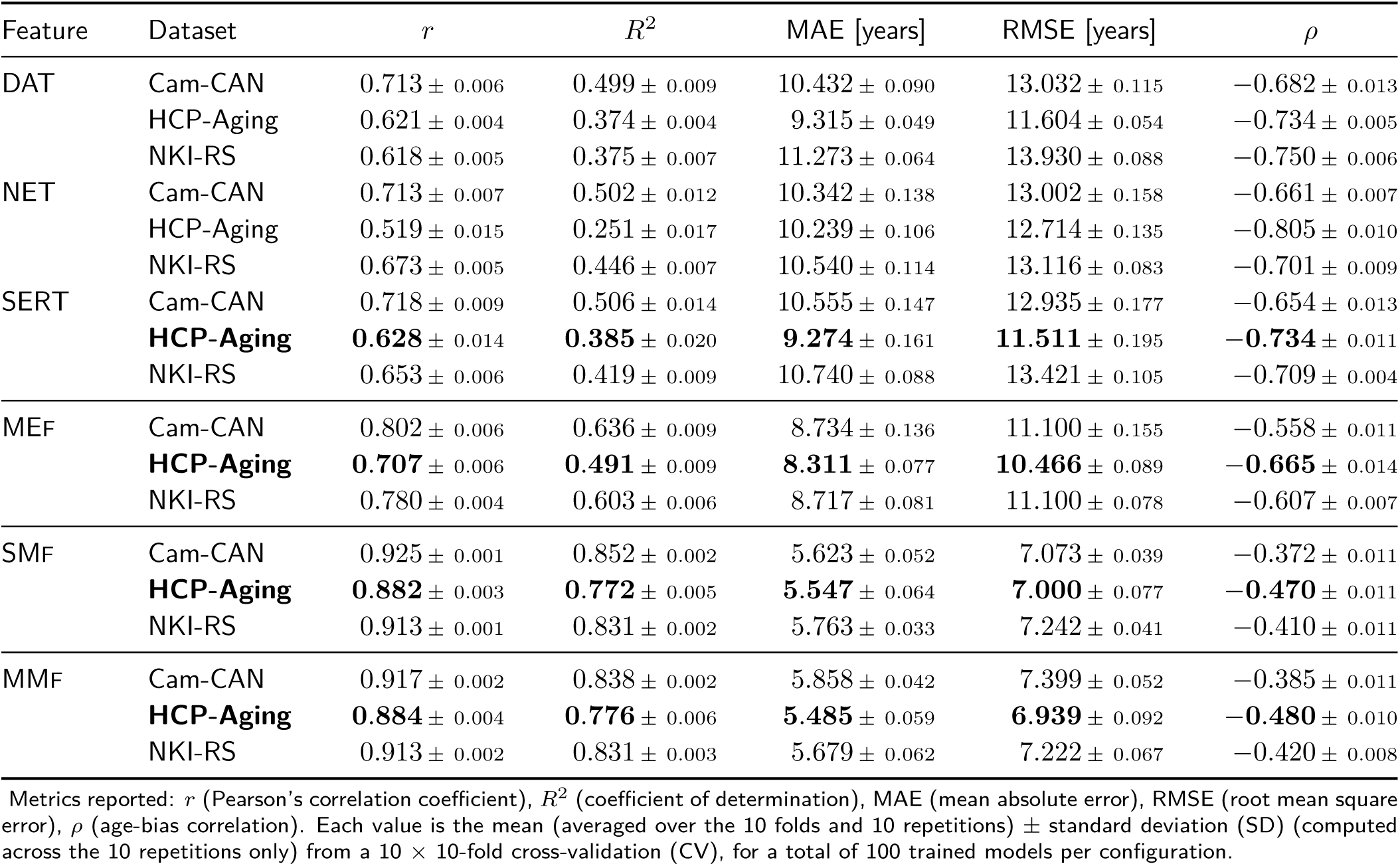
Single-site brain-age prediction performance across feature sets.

**Figure 3:**
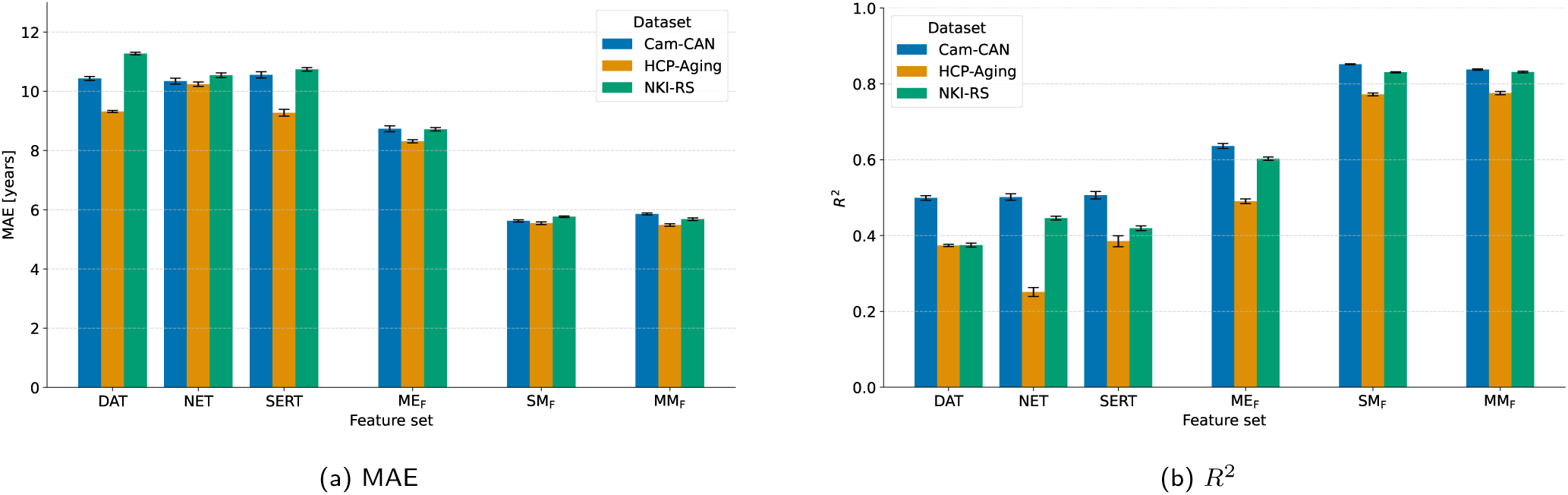
Comparison of model performance across feature sets (single transporter maps, molecular-enriched features (MEF), structural morphometric features (SMF), multimodal features (MMF)) between different datasets (Cam-CAN in blue, HCP-Aging in orange, NKI-RS in green): (a) mean absolute error (MAE) and (b) coefficient of determination (*R*^2^).

#### 3.3.2 Merged-cohort models: dataset harmonization is necessary

Performance for the merged cohort is shown in Table 5. For the raw (non-harmonized) data, single-map functional inputs produced MAEs of 11.00–11.27 years; MEF concatenation lowered the error to 9.67 years; and FS structural features achieved 5.33 years, whereas the MMF model did not improve that result (MAE = 5.59). Applying ComBat with empirical-Bayes pooling (CBE) raised DAT and MEF accuracy slightly but did not help NET or SERT, while structural performance remained stable. The independent-feature ComBat configuration (CBI) delivered the overall best figures, marginally improving SMF (achieving *R*^2^ of 0.86 with MAE = 5.26 years), while MMF achieved the next-lowest error (MAE=5.36 years). Thus, SMF harmonized with independent-feature ComBat represented the top multisite model, followed closely by the combined structural+functional feature set. The same pattern can be seen in Figure 4: harmonization improved performance overall, but did not reorder the hierarchy of the models.

#### 3.3.3 Common-parcellation analysis: parcellation mismatch explains the lost gains

The DKT-based analysis (Table 6) on the merged cohort (CBI harmonization) shows similar trends when reanalyzed on a common parcellation. Single-map functional inputs produced MAEs of 12.06–12.61 years, while the MEF concatenation reduced error to 10.45 years. SMF alone achieved an MAE of 6.02 years, and combining DKT structure with the three functional maps (MMF) lowered the error slightly further to 5.81 years with *R*^2^ of 0.83; compared to SMF, the absolute gain in performance is 0.21 years (3.5% improvement). These effects are visualized in Figure 5, which confirms that the joint DKT structural+functional model yields the lowest MAE and highest *R*^2^ within this parcellation framework.

**Table 5:**
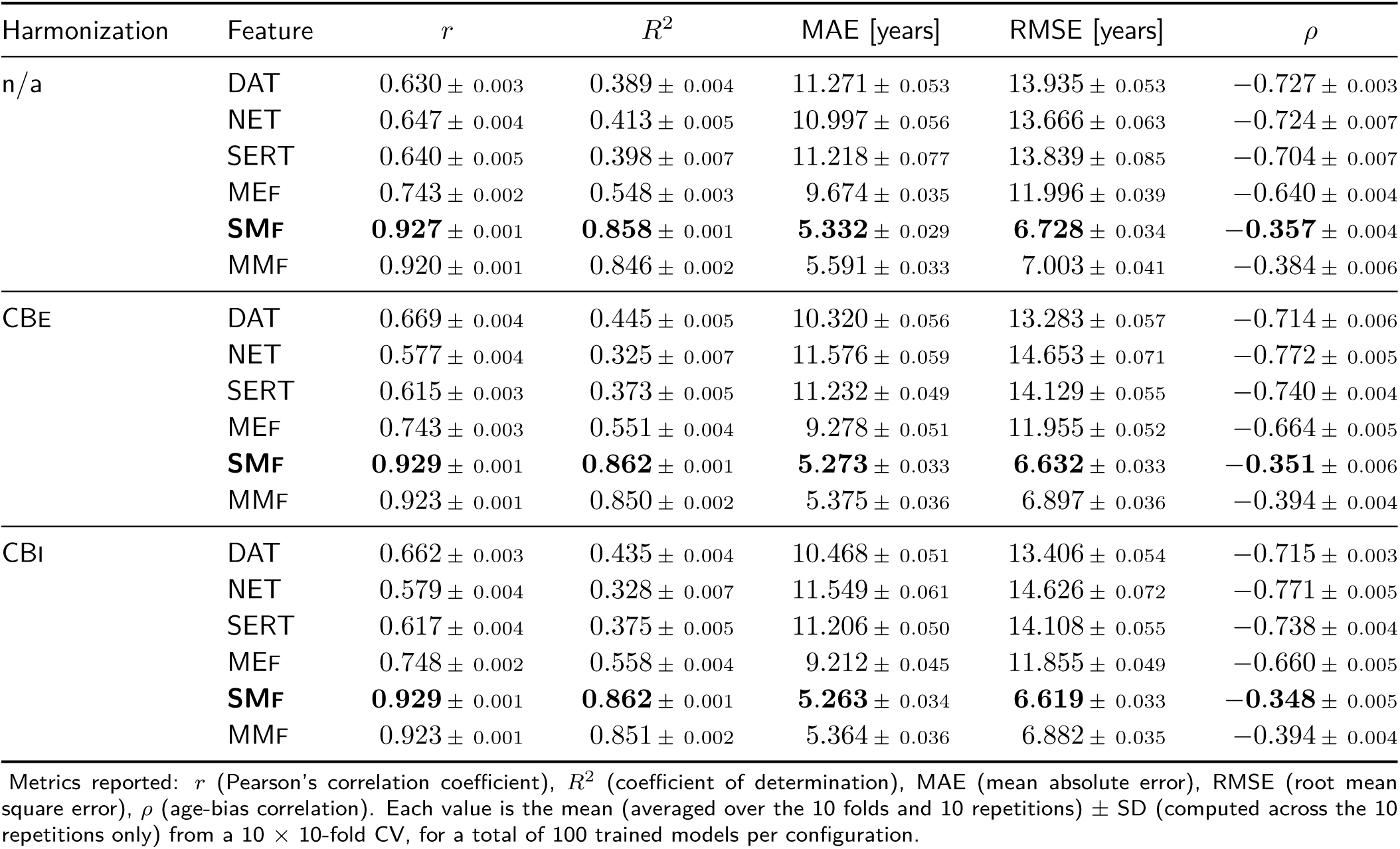
Multi-site brain-age prediction performance across harmonization procedures.

**Figure 4:**
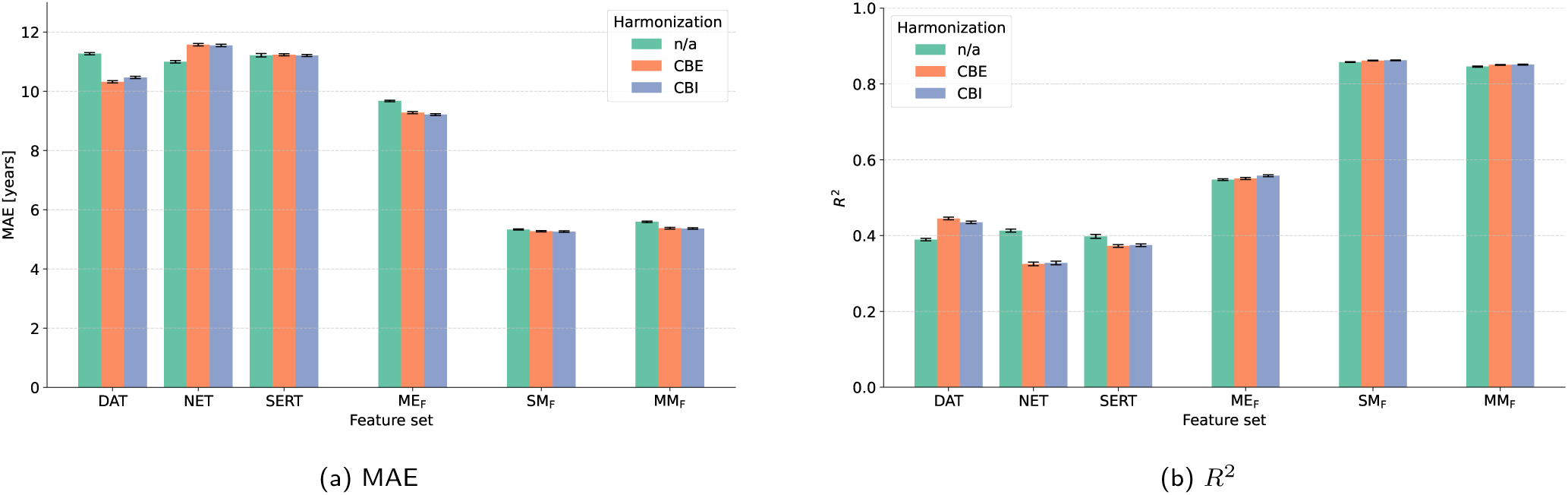
Comparison of model performance across feature sets for multi-site dataset and different harmonization strategies: (a) mean absolute error (MAE) and (b) coefficient of determination (*R*^2^).

**Table 6:** Multi-site brain-age prediction performance on common brain parcellation.

| Feature | $r$ | $R^2$ | MAE [years] | RMSE [years] | $\rho$ |
| --- | --- | --- | --- | --- | --- |
| DAT | $0.532 \pm 0.005$ | $0.267 \pm 0.006$ | $12.064 \pm 0.063$ | $15.272 \pm 0.065$ | $-0.774 \pm 0.006$ |
| NET | $0.477 \pm 0.004$ | $0.214 \pm 0.006$ | $12.610 \pm 0.039$ | $15.814 \pm 0.062$ | $-0.828 \pm 0.005$ |
| SERT | $0.517 \pm 0.004$ | $0.260 \pm 0.006$ | $12.281 \pm 0.043$ | $15.344 \pm 0.060$ | $-0.811 \pm 0.005$ |
| MEF | $0.657 \pm 0.003$ | $0.427 \pm 0.005$ | $10.453 \pm 0.039$ | $13.494 \pm 0.058$ | $-0.707 \pm 0.004$ |
| SMF | $0.908 \pm 0.001$ | $0.824 \pm 0.001$ | $6.017 \pm 0.031$ | $7.489 \pm 0.024$ | $-0.400 \pm 0.004$ |
| **MMF** | **$0.912 \pm 0.000$** | **$0.830 \pm 0.001$** | **$5.809 \pm 0.027$** | **$7.343 \pm 0.023$** | **$-0.398 \pm 0.002$** |

Metrics reported: $r$ (Pearson's correlation coefficient), $R^2$ (coefficient of determination), MAE (mean absolute error), RMSE (root mean square error), $\rho$ (age-bias correlation). Each value is the mean (averaged over the 10 folds and 10 repetitions) ± SD (computed across the 10 repetitions only) from a $10 \times 10$ -fold CV, for a total of 100 trained models per configuration.

**Figure 5:**
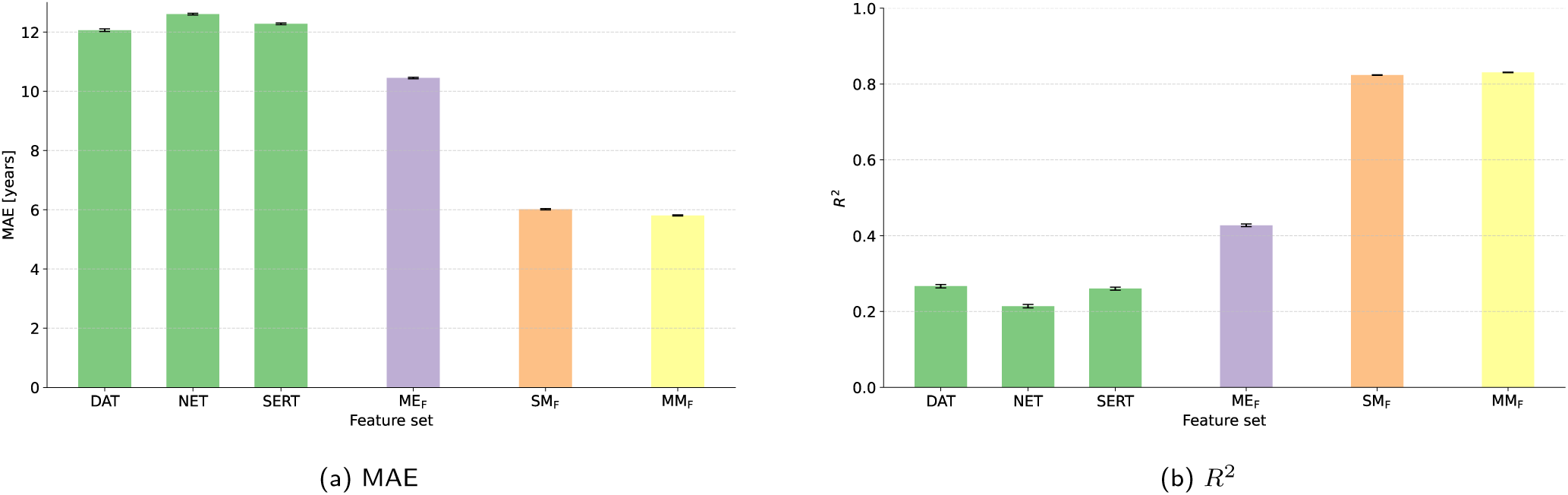
Comparison of model performance across feature sets on common brain parcellation: (a) mean absolute error (MAE) and (b) coefficient of determination (*R*^2^).

## 4 Discussion

We asked whether molecularly informed resting-state FC derived with REACT added predictive value to structural MRI for brain-age estimation. Using three adult cohorts (Cam–CAN, HCP–Aging, and NKI–RS), we trained SVR models on transporter-enriched maps (DAT, NET and SERT), FS morphometry, and their combination, and systematically assessed multi-site harmonization with ComBat. We then enforced a common parcellation to compute parcel-level features from both T1w and rs-fMRI modalities within the same anatomical node space, enabling direct cross-modality comparisons.

Across 2,120 adults, molecular-enriched FC was predictive of brain-age and, when added to morphometry, provided incremental value over a structure-only baseline. Within single cohorts, aggregating the three transmitter-weighted maps into one molecular-enriched FC block increased explained variance over any single transmitter map and, when added to FreeSurfer morphometry, yielded small, cohort-dependent gains. After merging datasets, harmonization was necessary to preserve the predictive value of molecular-enriched FC, with the independent-feature ComBat configuration (CBI) performing best and maintaining the multimodal advantage. Aligning modalities to a common atlas (DKT) restored a consistent improvement on the merged cohort, implicating parcellation mismatch as the principal reason gains appeared attenuated without alignment.

### 4.1 Brain-age with molecular-enriched FC

#### 4.1.1 Single-cohort models: cohort-dependent gains

Molecular-enriched FC alone predicted chronological age with moderate explained variance, and concatenating the three transmitter weightings reliably boosted *R*^2^ relative to any single map. Adding molecular-enriched FC to morphometry further improved performance in two cohorts (HCP–Aging and NKI–RS) while showing no gain in Cam–CAN. A minimal set of error metrics was consistent with this view: single-transmitter models had MAE on the order of 9–11 years, and concatenation typically reduced MAE by roughly 1–2 years per cohort, mirroring the rise in *R*^2^. The relative ranking of transmitter-specific maps varied by dataset. SERT/DAT performed slightly better than NET in Cam–CAN and HCP–Aging, whereas NET led in NKI–RS, suggesting cohort- and acquisition-dependent sensitivity of neuromodulatory circuits to aging. These findings indicate that transmitter-anchored connectivity captures unique age-related variance and yields a measurable, albeit cohort- and acquisition-dependent, incremental gain. Structural metrics remain a strong baseline, but molecular-enriched FC is itself predictive and can improve estimates.

#### 4.1.2 Merged-cohort models: harmonization is needed

Pooling Cam–CAN, HCP–Aging, and NKI–RS introduced site-related shifts that disproportionately affected functional features. Harmonization was essential to preserve and express the predictive value of molecular-enriched FC. Among tested strategies, CBI provided the best balance, recovering the strongest molecular-enriched FC block while maintaining high structural performance and the expected multimodal advantage. In line with the single-cohort trends, CBI modestly lowered error for molecular-enriched FC (e.g., from MAE 9.67 to 9.21 years for the concatenated block) and set a reference point for structure (5.26 years) without altering the qualitative hierarchy of feature sets. Taken together, these observations motivate a practical recommendation: deploy domain-specific pooling of harmonization parameters via CBI in multi-site brain-age pipelines, as it curbs scanner/site effects that otherwise degrade features.

#### 4.1.3 Common-parcellation analysis: atlas alignment restores molecular-enriched FC gains

Because functional gains appeared smaller after merging datasets, we examined whether parcellation mismatch masked true complementarity. Enforcing a shared cortical atlas (DKT) for both modalities clarified the picture: adding molecular-enriched FC to structure consistently improved prediction on the merged cohort, raising *R*^2^ from 0.824 to 0.830 with a parallel error reduction (from MAE 6.02 to 5.81 years). Thus, when node definitions are aligned, the incremental contribution of molecular-enriched FC is expressed reliably, and the earlier attenuation is best attributed to representational mismatch rather than weak functional signal.

In summary, these results converge on a coherent interpretation: molecular-enriched FC carries age-related signal that is not redundant with morphometry and, under routine multi-site constraints, can be translated into measurable improvements. Within single cohorts, concatenated transmitter weightings increase *R*^2^ and modestly reduce MAE; across cohorts, those gains are preserved when harmonization adopts CBI; and when modalities share the same parcellation, the multimodal advantage reappears consistently. In practice, treating molecular-enriched FC as a first-class feature block, paired with independent-feature ComBat and atlas alignment, yields brain-age models that retain mechanistic interpretability and, depending on cohort and alignment, match or surpass structure-only baselines.

### 4.2 Comparison with literature

Our findings are consistent with recent brain-age reports in both healthy and clinical cohorts. Using only T1w MRI, we achieved a MAE of roughly 5–6 years, matching the performance range described by Cole and Franke (2017). A study by Liem et al. (2017), which analyzed a slightly larger cohort of more than 2,300 adults and combined cortical thickness with whole-brain rs-FC features, reported a MAE of 4.29 years. In comparison, our best multimodal model, trained on about 2,100 participants and integrating FreeSurfer morphometry with molecular-enriched FC, reached a MAE of 5.36 years. The small gap likely reflects differences in sample composition and the added variability introduced by pooling data across scanners. Importantly, our SMF model attained an *R*^2^ near 0.85, a level comparable to those single-site studies.

Liem et al. (2017) also observed that adding FC to structural features improved accuracy. In our data, structural metrics were already highly predictive, so the molecular-enriched FC features provided only a modest additional benefit; nonetheless, they did lower the residual error and increased explained variance, confirming that age-related changes in functional networks convey information not captured by anatomy alone. This discrepancy may be due to differences in functional feature richness: Liem et al. (2017) used full resting-state connectivity matrices, which encompass aging effects on diverse networks, whereas we focused on specific neurotransmitter-enriched connections. It is plausible that our functional features, being constrained to DAT-, NET-, and SERT-related circuits, captured a narrower slice of age-related functional change, thus emphasizing neurochemical network effects but potentially missing other widespread changes (e.g., in default-mode or visual networks) that whole-brain FC approaches exploit. Future studies could explore augmenting our molecular connectivity features with broader functional measures to see whether multimodal combinations can surpass structure-only models, as reported by Liem et al. (2017) and others.

Recent work has indeed demonstrated that resting-state FC alone can be a potent predictor of age. Bi et al. (2024) used the Cam–CAN dataset and reported that both movie-watching and resting-state FC yielded high correlation with age (*R*^2^=0.74). This accuracy is comparable to, and in some cases exceeds, the performance of structural MRI models. Our results temper this optimistic view by showing that, in a direct comparison within the same individuals, structural features still provided substantially higher *R*^2^ (up to 0.85) than any single functional feature (*R*^2^=0.51 at best), and even the multi-network FC model (MEF) reached only *R*^2^=0.64 (Table 4). It is important to note that Bi et al. (2024) and similar studies typically employ many more FC features (e.g., thousands of connections) and often use advanced machine-learning or deep-learning models that can capture non-linear aging patterns in connectivity. In contrast, our approach emphasizes interpretability by using predefined molecular circuits and a relatively simple SVR model. The trade-off is a lower ceiling on FC-based accuracy, but with the benefit of linking age effects to specific neurochemical systems. Encouragingly, our MEF model’s MAEs (8.31–8.72 years single-site, 9.21 years multi-site) are on par with other moderate-sized FC studies (Gonneaud et al., 2021; Millar et al., 2022; Zhai & Li, 2019), and with further refinement these numbers could improve.

Our work also connects with emerging findings in pathological aging. Millar et al. (2022) examined brain-age in preclinical and symptomatic Alzheimer’s disease (AD) using functional and structural MRI. They found that an FC-only age model detected aberrant brain aging in asymptomatic individuals at risk for AD, even when structural MRI appeared normal. In our healthy dataset we do not directly test disease effects, but the strong performance of structural features implies they capture the lion’s share of normative aging variance, whereas FC may capture subtler, system-specific changes that could represent early signs of neurodegeneration before structural atrophy becomes evident. Intriguingly, Millar et al. (2022) reported that while both functional and structural brain-age gap metrics were elevated in cognitively impaired AD patients, only the FC-based brain age showed a paradoxical decrease (younger-appearing brains) in the preclinical stage, possibly reflecting transient network reorganization. Such complex, bi-phasic effects underscore that FC and structural measures are complementary: FC might be more sensitive to early or compensatory changes, whereas structural changes correlate more linearly with cumulative damage.

Although we did not find a major boost in accuracy from adding FC in healthy adults, it is likely that in a clinical context the functional features would add important diagnostic value, e.g., flagging individuals whose brain connectivity appears “older” or “younger” than expected for their age in ways that structural MRI cannot. In the literature, several multimodal brain-age studies (e.g., Liem et al., 2017; Millar et al., 2023) conclude that integrating across modalities yields the most robust predictors and better captures cognitive or pathological deviations. To our knowledge, this is the first large-scale, multi-site brain-age investigation to incorporate molecular-enriched FC, adding a molecularly informed functional dimension to brain-age estimation. Future comparisons with other multimodal frameworks will clarify optimal strategies for fusing anatomical, functional, and molecular imaging data.

### 4.3 Limitations

This study nonetheless has a few noteworthy limitations. First, although we used a non-linear support-vector regression with a radial-basis-function kernel, SVR remains constrained by pair-wise similarity structure and a relatively small hyper-parameter set, which can limit its ability to model complex feature interactions. In line with this, our prior benchmarking across kernel-based and ensemble methods showed that model choice and performance depend strongly on data distribution and harmonization: gradient-boosted trees (XGB) were most robust on non-harmonized data, whereas SVR achieved the best accuracy after ComBat (Pinamonti & Sammassimo et al., 2025). These results suggest that no single algorithm is uniformly optimal and motivate exploration of larger training sets and higher-capacity models (e.g., ensembles or deep learning), while acknowledging trade-offs in interpretability and over-fitting risk. Second, we extracted atlas-level features and compared results across two different parcellations. Alternative parcellations, or voxel-wise, surface-based, or graph-theoretic representations, could reveal finer-grained or topological aging effects, particularly in functional connectivity. Third, our multi-site harmonization relied on ComBat. While overall performance improved, effects were uneven and transmitter-dependent: NET deteriorated under both harmonization variants and SERT remained essentially unchanged. These patterns suggest possible over-correction for some features or residual entanglement between site and age effects; future work could explore site-invariant representation learning or hierarchical models. Fourth, the study is cross-sectional. Without longitudinal follow-up we cannot determine whether participants with higher brain-age gaps actually age faster. Longitudinal validation and uncertainty quantification are essential for clinical deployment, where single-subject decisions require tighter error bounds than the 5-years MAE achieved here. Fifth, molecular enrichment was based on population-average PET/SPECT templates for only three monoaminergic transporters. Inter-individual variability in receptor density and aging effects in other neurotransmitter systems were not modeled. Incorporating subject-specific molecular imaging or additional receptor atlases could enhance sensitivity. Finally, our analytic sample comprised healthy adults from Cam-CAN (UK), HCP-Aging (USA), and NKI-RS (USA), screened as cognitively normal according to each cohort’s original protocols. Generalizability to other demographic, socioeconomic, or clinical populations remains to be established and should be tested in more diverse, longitudinal cohorts.

## 5 Conclusion

Our goal was to advance brain-age modeling by pairing established morphometric predictors with novel, molecular-enriched FC features and to test their performance across multiple imaging sites. Consistent with our hypothesis, structural measures yielded the highest standalone accuracy, with MAE in the five-to-six-year range and excellent generalizability across scanners.

Interestingly, the inclusion of neurotransmitter-weighted FC maps into a brain-age model explained a significant amount of brain age variance that structure alone did not capture. When DAT, NET, and SERT-enriched networks were added to the model, prediction error decreased, indicating that functional alterations within monoaminergic circuits contribute complementary aging-related information beyond structural anatomy. These gains, though modest in absolute terms, were consistent across atlases and remained after ComBat harmonization, underscoring that neuromodulatory connectivity holds independent predictive value beyond macroscopic atrophy.

The findings therefore highlight two complementary facets of cerebral aging: widespread tissue loss, which drives the bulk of structural prediction, and systematic alterations in neurochemical functional networks, which refine, and potentially re-interpret, brain-age estimates. Future work should deepen this molecular perspective by integrating a broader repertoire of receptor systems, employing individual PET/SPECT maps where available, and examining longitudinal or clinical cohorts in whom neuromodulatory dysfunction may emerge before overt atrophy. Such efforts could transform molecular-enriched FC from a modest performance booster into a sensitive early marker that enriches the biological meaning and clinical utility of brain-age metrics.

## Data and Code Availability

All neuroimaging data used in this study are publicly accessible. T1w and rs-fMRI scans from CamCAN are available at https://camcan-archive.mrc-cbu.cam.ac.uk/dataaccess/ (Shafto et al., 2014; Taylor et al., 2017). HCP-Aging data can be downloaded from https://www.humanconnectome.org/study/ hcp-lifespan-aging (Bookheimer et al., 2019; Harms et al., 2018). NKI-RS data are provided at https://fcon_1000.projects.nitrc.org/indi/enhanced/ (Nooner et al., 2012; Tobe et al., 2022).

Preprocessing pipelines relied exclusively on publicly available BIDS-Apps: functional MRI Preprocessing (fMRIPrep) (v23.2.2, https://fmriprep.org/en/23.2.2/), XCP-D (v0.7.4, https://xcp-d.readthedocs.io/en/0.7.4/), and MRIQC (v24.1.0, https://mriqc.readthedocs.io/en/latest/) or common neuroimaging tools as FSL (v7.3.2, https://fsl.fmrib.ox.ac.uk/fsl/docs/). REACT was implemented with the open-source react-fmri package, available at https://github.com/ottaviadipasquale/react-fmri. Custom scripts for brain-age modeling (SVR) and feature harmonization with ComBat were written for local execution and are available upon reasonable request from the corresponding author under a standard data-use agreement.

## Author Contributions

**Marco Pinamonti:** Conceptualization; Methodology; Data curation; Software; Formal analysis; Writing - original draft. **Manuela Moretto:** Methodology; Data curation; Supervision; Writing - review & editing. **Valentina Sammassimo:** Methodology; Software; Validation; Formal analysis; Writing - review. **Marco Castellaro:** Resources; Software; Data curation; Writing - review. **Mattia Veronese:** Conceptualization; Supervision; Funding acquisition; Writing - review & editing.

All authors have read and approved the final manuscript.

## Funding

This research was funded by the Ministry of University and Research within the Complementary National Plan PNC-I.1, “Research initiatives for innovative technologies and pathways in the health and welfare sector, D.D. 931 of 06/06/2022, PNC0000002 DARE - Digital Lifelong Prevention CUP: B53C22006440001.”

## Declaration of Competing Interests

The authors declare that they have no competing interests.

## Acknowledgments

We acknowledge the use of data from three public databases.

Data collection and sharing for this project were provided by the Cambridge Centre for Ageing and Neuroscience. Cambridge Centre for Ageing and Neuroscience funding was supported by the UK Biotechnology and Biological Sciences Research Council (grant number BB/H008217/1), together with support from the UK Medical Research Council and the University of Cambridge, UK.

Additionally, data from the HCP-Aging 2.0 Release were utilized in this research. Research reported in this publication was supported by the National Institute on Aging of the National Institutes of Health under Award Number U01AG052564 and by funds provided by the McDonnell Center for Systems Neuroscience at Washington University in St. Louis. The Lifespan Human Connectome Project in Aging 2.0 Release data used in this report came from DOI: 10.15154/1520707.

We also acknowledge the Enhanced Nathan Kline Institute–Rockland Sample for providing open-access neuroimaging data; data collection and sharing for the Enhanced Nathan Kline Institute–Rockland Sample were supported by National Institute of Mental Health grant R01MH094639-01 and by additional funding from the New York State Office of Mental Health, the Research Foundation for Mental Hygiene, the Child Mind Institute (1FDN2012-1), the Center for the Developing Brain at the Child Mind Institute, National Institute of Mental Health grants R01MH081218, R01MH083246, and R21MH084126, the NKI Center for Advanced Brain Imaging, the Brain Research Foundation, and the Stavros Niarchos Foundation.

## Supplementary Material Material

### Subjects excluded

#### Quality control of raw images

Automated MRIQC screening flagged a subset of scans with poor signal-to-noise ratio, motion artifacts, or other anomalies; those participants were removed before preprocessing. In total, 61 individuals were excluded: 51 because of structural issues and 10 because of functional issues. The dataset-specific counts are listed in Supplementary Table S1.

**Table S1:** Participants removed after MRIQC.

| Dataset | Structural (T1w) | Functional (rs-fMRI) | Total |
| --- | --- | --- | --- |
| Cam-CAN | 3 | 2 | 5 |
| HCP-Aging | 25 | 8 | 33 |
| NKI-RS | 23 | 0 | 23 |
| **Total** | **51** | **10** | **61** |

#### Preprocessing outputs

During structural and functional MRI preprocessing, a small additional fraction of scans failed because of unrecoverable software errors or excessive motion. Specifically, *fMRIPrep* terminated early for five HCP-Aging participants, whereas no such failures occurred in Cam-CAN or NKI-RS. Subsequent denoising with XCP-D led to the exclusion of participants whose residual time series contained more than one-third of volumes with framewise displacement exceeding 0.5 mm. Finally, one subject was excluded from Cam-CAN sample because react-fmri did not run successfully on it. Table S2 details these counts: in total, 165 participants were removed at the preprocessing stage, leaving 2128 individuals for downstream analyses. This number lowers to 2120 after the exclusion of HCP-Aging participants over 90 years old: this final sample constitutes the dataset used for this study.

#### Molecular-enriched functional connectivity

Only one run failed (subject from Cam-CAN) and thus that participant was excluded from the analysis.

**Table S2:** Participants excluded during preprocessing.

| Dataset | fMRIPrep | XCP-D | REACT | Total |
| --- | --- | --- | --- | --- |
| Cam-CAN | 0 | 20 | 1 | 21 |
| HCP-Aging | 5 | 79 | 0 | 84 |
| NKI-RS | 0 | 60 | 0 | 60 |
| Total | 5 | 159 | 1 | 165 |

## Footnotes

1 https://github.com/bids-standard/bids-validator, https://bids-validator.readthedocs.io/en/latest/

2 https://github.com/nipreps/mriqc, https://mriqc.readthedocs.io/

## Notes

### Competing Interest Statement

The authors have declared no competing interest.

